# Sex-stratified immune patterns across the canine lifespan and short-term responses to geroprotective interventions

**DOI:** 10.64898/2026.09.09.750505

**Authors:** Meiling Lai, Fengge Xu, Yingxia Xu, Albert Mironenkov, Yi Jin, Shengxiang Zhang, Jianhong Pan, Ming Li, Baichuan Deng, Jian-Kang Zhu, Yu-Xuan Lyu

**Author notes:** Meiling Lai, Fengge Xu and Yingxia Xu contributed equally to this work. Corresponding authors (Yu-Xuan Lyu), (Jian-Kang Zhu), (Baichuan Deng), (Ming Li).

## Abstract

Canine models are invaluable for translational geroscience, but natural sex-associated immune aging remains underinvestigated. Here, we characterize the age-associated hematologic and serum cytokine profiles in a controlled cross-sectional cohort of 80 intact laboratory Beagles spanning 1 to 11 years of age. Canine immune aging is heterogeneous and non-linear. While several absolute leukocyte counts declined with age before rising again in geriatric dogs, serum cytokines exhibited distinct, sex-stratified patterns. Although more cytokines varied significantly with age in males than in females, formal age-by-sex interactions did not remain significant after false discovery rate correction. To explore short-term geroprotective responses, we evaluated 24 young beagles following a 90-day intervention with rapamycin, canagliflozin, or dietary restriction. Rapamycin was associated with the broadest endpoint cytokine response. Canagliflozin showed narrower cytokine differences alongside descriptive body-weight reduction, whereas dietary restriction reduced body weight without altering measured immune endpoints. Given the small group sizes and endpoint-only cytokine measurements, these intervention findings remain exploratory and hypothesis-generating. Together, our findings describe stage-specific hematologic changes and sex-stratified cytokine patterns across the canine lifespan and provide a basis for future longitudinal studies.

## Introduction

Aging is a gradual functional decline across organ systems, and it is the major risk factor for numerous life-threatening chronic diseases (Kennedy et al. 2014; López-Otín et al. 2023). A central aim of geroscience is to understand the shared mechanisms behind this decline and to develop interventions that delay it (Kennedy et al. 2014). Many core aging pathways have been discovered utilizing short-lived models, such as the yeast, worms, fruit flies, and mice (López-Otín et al. 2023). However, these models differ from people in lifespan, body size, living environment, and disease spectrum, which limits direct translation (Hoffman et al. 2018; Ruple et al. 2022). Domestic dogs offer a complementary model as they share the human environment, receive veterinary care, and develop naturally occurring age-related diseases (Creevy et al. 2022; Hoffman et al. 2018; Ruple et al. 2022). They are also shorter-lived than people, so adult and late-life functional changes can be followed within a feasible time frame (Kaeberlein et al. 2016; Ruple et al. 2022). Together, these features place domestic dogs as a premier translational model for human aging research (Creevy et al. 2022; Kaeberlein et al. 2016).

This translational potential has led to a growing ecosystem of canine aging studies and early intervention trials. The Dog Aging Project follows large numbers of companion dogs to identify biological, environmental, and clinical determinants of aging, and it includes an interventional rapamycin arm (Coleman et al. 2025; Creevy et al. 2022; Kaeberlein et al. 2016). Other cohorts, including family-dog studies, the Golden Retriever Lifetime Study, and retired sled-dog programs, address cognition, disease risk, and standardized aging trajectories (Fleyshman et al. 2021; Labadie et al. 2022; Piotti et al. 2018; Urfer et al. 2021). Together, these efforts show that canine aging research is moving from descriptive gerontology toward cohort-based, mechanistic, and interventional geroscience. However, the precise trajectories of immune and hematologic aging remain incompletely characterized, especially regarding natural sex differences under standardized experimental conditions.

Aging trajectories are profoundly shaped by sex — a dimension that remains largely underexplored in aging research. Males and females often differ in lifespan, in the pace of functional decline, and in the risk of many age-related diseases (Austad and Fischer 2016; Hägg and Jylhävä 2021). Yet, many studies using animal models still include only one sex, or report results without stratifying by sex (Klein and Flanagan 2016). This practice can hide sex-specific effects and weaken translation. Therefore, including sex as a biological variable is indispensable for rigorous geroscience and for evaluating potential differences in geroprotective responses.

The immune system is a sensitive readout of aging, and it is also strongly shaped by sex. In humans, men and women exhibit distinct baseline inflammatory tones and divergent rates of immunosenescence and inflammaging, driven in part by the sex chromosomes and differential decline of sex hormones (Ferrucci and Fabbri 2018; Franceschi et al. 2000; Franceschi et al. 2018; Gubbels Bupp 2015; Klein and Flanagan 2016). Multi-omic profiling shows that human immune aging follows sex-specific trajectories, with different genes, cell subsets, and cytokine patterns changing with age in each sex (Márquez et al. 2020). Similar immune-aging features have been described in dogs. Companion dog studies have linked older age to lower lymphocyte counts and shifts in inflammatory markers (McMahon et al. 2025; Schmid et al. 2024). Cross-sectional and longitudinal work reports age-related changes in leukocytes, immunoglobulins, and cytokines such as IL-6, IL-8, and TNF-α (Alexander et al. 2018; Jiménez 2023; McMahon et al. 2025; Schmid et al. 2024). Routine hematologic and biochemical measures also change with age, and they are now used to build canine biological age markers (Herzig et al. 2025; Radakovich et al. 2017; Vajdovich et al. 1997; Zemko et al. 2025).

Despite this progress, mapping natural canine immune aging is still confounded by several major variables. First, many studies use companion dogs, where breed, body size, diet, environment, disease, and neuter status all add substantial variation (Hart et al. 2020; Jiménez 2023; Sundburg et al. 2016). Crucially, the widespread practice of gonadectomy in companion dogs fundamentally alters sex hormone axes. Because androgens and estrogens regulate immune-cell function and systemic inflammation, gonadectomy may alter or mask natural age-associated immune patterns. Profiling immune aging in standardized, intact canine cohorts can therefore help assess sex-stratified patterns without the additional counfounding of neuter status. Second, many studies use broad age categories or assume a linear change with age. This can hide stage-specific or non-linear patterns. In humans, by contrast, plasma proteins and multi-omic profiles change in waves rather than along a straight line (Lehallier et al. 2019; Shen et al. 2024). Whether canine immune markers behave in a similar way is unclear. Third, it remains uncertain whether blood cell counts and serum cytokines change together or follow different axes of immune aging. Systematic profiling of both blood and cytokine measures in a standardized and sex-balanced dog cohort remains limited.

A parallel aim of geroscience is to test candidate geroprotective interventions, several of which exhibit notable sex biases (Kennedy et al. 2014). Three candidates are especially relevant. Rapamycin inhibits mTOR and extends lifespan in various models including mice (Harrison et al. 2009). Canagliflozin, a sodium-glucose cotransporter 2 inhibitor, extends median lifespan in male but not female mice, and it alters metabolism and neuroinflammation in a sex-dependent manner (Jayarathne et al. 2022; Miller et al. 2020). This sex-biased effect makes canagliflozin a powerful probe for sex differences in intervention response. Long-term dietary restriction extends lifespan and delays disease in several species, including dogs (Kealy et al. 2002; Lawler et al. 2008), and shorter-term calorie restriction reshapes immune function in humans (Spadaro et al. 2022). While canine intervention research is growing, it remains limited in scope. Rapamycin has been tested in companion dogs, mainly for safety and cardiac endpoints, and a larger trial is still underway (Barnett et al. 2023; Coleman et al. 2025; Urfer et al. 2017). Other work has paired transcriptomic aging signatures with anti-aging interventions in dogs (Zeng et al. 2024). Yet, few studies compare the early effects of different candidate interventions within one framework. Standardized Beagle cohorts offer a complementary setting, where interventions can be compared under a more consistent genetic background, diet, housing, and management. Furthermore, short-term treatments in young dogs can describe early immune and hematologic responses, generating hypotheses for later midlife and late-life studies.

Here, we address these translational gaps in a standardized cohort of intact laboratory Beagles. We first investigated complete blood count (CBC) parameters and serum cytokine profiles across five age groups, from young adult to geriatric dogs, and analyzed these data separately in males and females. We then performed a 90-day intervention in young intact beagles to compare rapamycin, canagliflozin, and dietary restriction, measuring body weight, leukocyte counts, and serum cytokines. Together, these data provide a sex-stratified description of canine immune aging and exploratory data on short-term responses to candidate interventions.

## Methods

### Cross-sectional aging study design

The cross-sectional aging study included 80 intact laboratory Beagles, comprising 40 males and 40 females. Before enrollment, all dogs underwent physical examination, and animals with overt organ-system disease were excluded. Dogs were stratified by chronological age into five predefined age groups: junior dogs aged 1.3-1.5 years (n = 12; 6 males and 6 females), adult dogs aged 3.3-4.3 years (n = 12; 6 males and 6 females), middle-aged dogs aged 6.2-6.6 years (n = 12; 6 males and 6 females), senior dogs aged 8.7-9.3 years (n = 24; 12 males and 12 females), and geriatric dogs aged 10.1-10.5 years (n = 20; 10 males and 10 females). Dogs in this cohort were fed 200 g/day of a standard commercial canine maintenance diet. Estrous stage was not recorded or experimentally controlled at blood collection. None of the female dogs was pregnant, lactating or being used for breeding at sampling.

### Intervention study design

The intervention study included 24 young laboratory Beagles, comprising 12 males and 12 females, and lasted for 3 months. Before enrollment, all dogs underwent physical examination, and animals with overt organ-system disease were excluded. Eligible dogs were allocated to four intervention groups using an age-, sex-, and body weight-balanced stratification strategy: control, rapamycin, canagliflozin, and dietary restriction. The control group included dogs aged 0.8-1.3 years (n = 6; 3 males and 3 females), which were fed 250 g/day and received empty capsules. The rapamycin group included dogs aged 0.9-1.1 years (n = 6; 3 males and 3 females), which were fed 250 g/day and received oral rapamycin capsules at 0.50 mg/kg/week, administered as 0.25 mg/kg per dose on Monday and Thursday mornings. The canagliflozin group included dogs aged 0.9-1.3 years (n = 6; 3 males and 3 females), which were fed 250 g/day and received oral canagliflozin capsules at 6 mg/kg/day once daily. The dietary-restriction group included dogs aged 0.8-1.3 years (n = 6; 3 males and 3 females), which were fed 185 g/day and received no pharmacological intervention or empty capsules. According to the manufacturer’s specifications, the diet had a metabolizable energy density of 3.7 kcal/g. Thus, the daily energy allowances were 925 kcal for the control group and approximately 685 kcal for the dietary-restriction group. The 185-g allowance was 26% lower by mass and calculated energy intake than the 250 g/day provided to the age- and activity-matched concurrent control group.

### Animal housing and ethics

All dogs were housed at the laboratory animal facility of Anling Biomedical (Shenzhen) Co., Ltd., which was maintained at 16-28 °C and 30-70% relative humidity under a 12-h light/12-h dark cycle. Unless otherwise specified, the dogs were fed a standard commercial canine maintenance diet (Canine Maintenance Formula Feed, product no. 5213; Keao Xieli (Tianjin) Feed Co., Ltd., Tianjin, China). Water was provided ad libitum. All animal procedures were approved by the Institutional Animal Care and Use Committee of Anling Biomedical (Shenzhen) Co., Ltd. (approval no. 2025-02-ALSHZH-21). The procedures complied with applicable institutional and national guidelines and the NIH Guide for the Care and Use of Laboratory Animals. The study is reported in accordance with the ARRIVE guidelines.

### Preparation of intervention capsules

Individual treatment doses were calculated once using each dog’s body weight measured before the intervention and remained fixed throughout the 90-day intervention. All treatment capsules were prepared before the start of the intervention.

Rapamycin (sirolimus) powder was provided by North China Pharmaceutical Co., Ltd. (Shijiazhuang, China). For each dog, rapamycin powder was weighed at 0.25 mg/kg using a microanalytical balance (XPR36DR/A; Mettler-Toledo GmbH, Greifensee, Switzerland) and transferred into size 1 enteric-coated empty hard gelatin capsules (Shanghai Redstar Capsules Co., Ltd., Shanghai, China).

Canagliflozin tablets (Invokana®, 100 mg per tablet; Janssen-Cilag S.p.A., Latina, Italy) were cut into fragments. For each dog, the required fragment mass to achieve a dose of 6 mg/kg was calculated based on the labeled drug content and the measured mass of one intact tablet. The calculated amount was weighed and placed into size 1 enteric-coated hard gelatin capsules (Shanghai Redstar Capsules Co., Ltd., Shanghai, China).

### Complete blood count analysis

Fasting peripheral blood (2 mL) was collected from each dog by forelimb venipuncture into K_2_EDTA anticoagulant tubes. Samples were analyzed immediately after collection, and all complete blood count (CBC) measurements were completed within 4 h. CBC testing was performed at Anling Biomedical (Shenzhen) Co., Ltd. Measurements were obtained using an automated hematology analyzer (XN-2000V; Sysmex Medical Electronics [Shanghai] Co., Ltd., Shanghai, China). CBC parameters included leukocyte, erythrocyte, platelet, and reticulocyte indices.

### Serum cytokine measurement

For serum cytokine analysis, all dogs underwent an overnight fast, and blood was collected between 09:00 and 11:00. A 5-mL peripheral blood sample was collected from each dog by forelimb venipuncture into vacuum tubes without anticoagulant. Blood samples were allowed to clot at room temperature for 2 h and then centrifuged at 3200 rpm for 15 min. The serum supernatant was collected, aliquoted at 250 μL per vial, and stored at -80 °C until analysis.

Serum cytokines were measured in the selected serum samples using the Quantibody Canine Cytokine Antibody Array 40 Kit (QAC-CAA-40; RayBiotech, Inc., Norcross, GA, USA) according to the manufacturer’s instructions. For the intervention study, cytokine profiling was performed at the 3-month endpoint. Each cytokine was measured using four technical replicate spots per sample, and the coefficient of variation among replicate spots was <15%. After background subtraction, the four replicate signals were averaged. Samples from the cross-sectional and intervention cohorts were distributed across different slides so that no study group was concentrated on a single slide. Q-Analyzer software performed intra-slide and inter-slide normalization against the fixed biotinylated positive-control spots on each array. The software then converted the normalized fluorescence signals to absolute cytokine concentrations using analyte-specific standard curves. The resulting concentrations (pg/mL) were used for downstream analysis. No cytokines were excluded because of poor technical performance.

### Statistical analysis

All statistical analyses and data visualization were performed using Python 3.12, R 4.3.2, and GraphPad Prism 10.4.1. Sample sizes, data summaries, and panel-specific tests are specified in the corresponding figure legends. Unless stated otherwise, tests were two-sided, with *P* < 0.05 or adjusted *P* value < 0.05 considered statistically significant.

The cross-sectional CBC analysis included 79 dogs because CBC data were unavailable for one dog. Two-way ANOVA was used to assess the effects of age group, sex, and their interaction on 17 CBC parameters. For each model term, *P* values across the 17 parameters were adjusted separately using the Benjamini-Hochberg procedure. Spearman rank correlations were used to assess associations between continuous age and five leukocyte outcomes (WBC, NEUT#, LYMPH#, MONO# and BASO#). The five corresponding *P* values were adjusted together using the Benjamini-Hochberg procedure. The complete nominal and adjusted *P* values are provided in Supplementary Table 4. Because the dogs occupied narrow predefined age bins separated by substantial gaps, age-group ANOVA was the primary approach, and Spearman correlation was used as a complementary test of overall monotonic trends.

Cytokine concentrations were analyzed as log2(pg/mL + 1). The cross-sectional cytokine analysis used a balanced dataset of five dogs per age-by-sex stratum across four age groups (n = 40). For each of the 40 cytokines, two-way ANOVA was used to assess the effects of age group, sex and the age-group-by-sex interaction. For each model term, *P* values were adjusted separately across the 40 cytokines using the Benjamini-Hochberg procedure, and partial ηp² was calculated as the effect-size measure. To characterize age-associated cytokine patterns within each sex, age-group effects were also assessed separately in males and females using one-way ANOVA. *P* values were adjusted across 40 cytokines within each sex using the Benjamini-Hochberg procedure.

Body-weight trajectories were summarized descriptively during the 3-month intervention. At the 3-month endpoint, each intervention group was compared with the concurrent control group using two-sided unpaired Welch’s t-tests for CBC parameters. The 12 *P* values were adjusted together using the Benjamini-Hochberg procedure. The complete nominal and adjusted *P* values are provided in Supplementary Table 5.

## Results

### Cross-sectional and intervention cohorts

To characterize age-associated blood phenotypes, we analysed a cross-sectional cohort of 80 intact laboratory Beagles distributed across five age groups (Fig. 1). Complete blood count (CBC) data were available for 79 dogs. Because array capacity was limited, serum cytokines were profiled in a balanced subset of 40 dogs. This subset comprised five males and five females from each of the junior, middle-aged, senior and geriatric groups. The adult group was not included in the cytokine analysis.

**Fig. 1.**
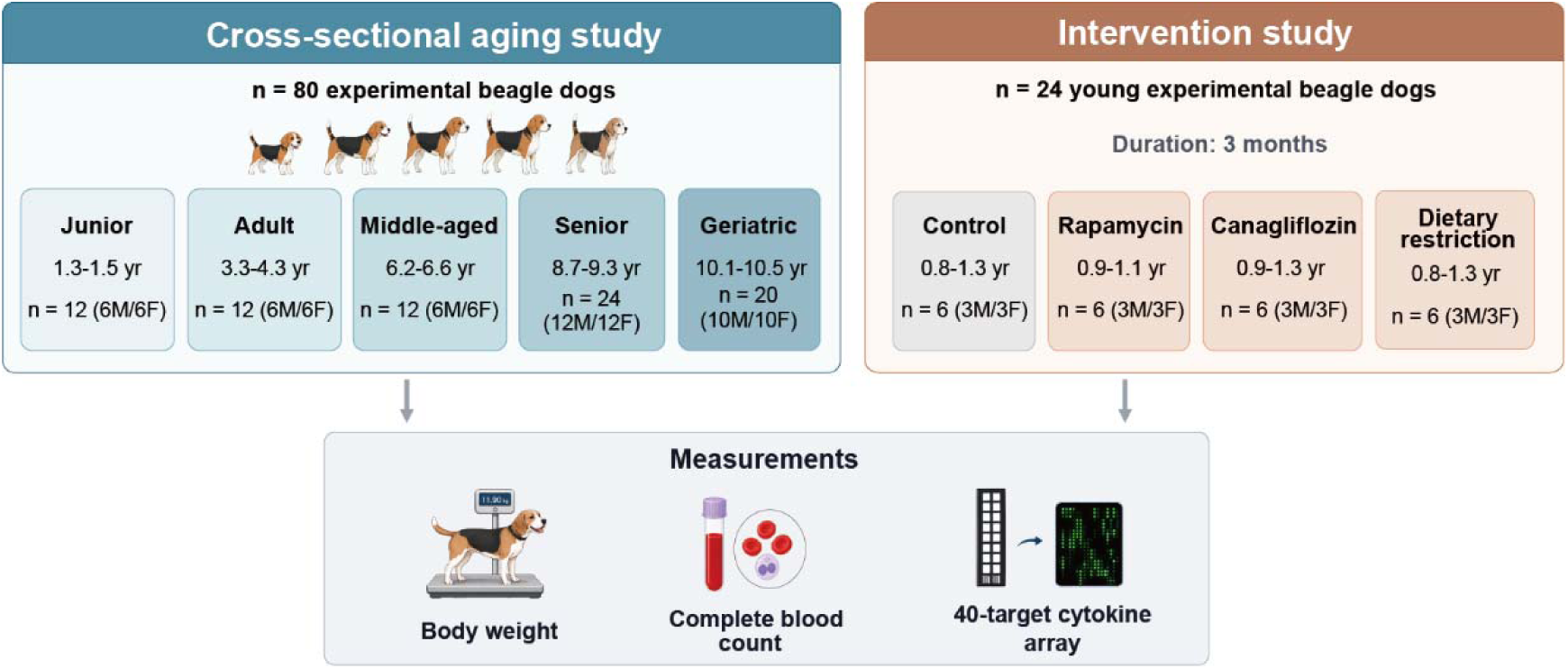
Study design of the cross-sectional aging and intervention studies. The cross-sectional study comprised 80 laboratory Beagles: junior, adult, middle-aged, senior and geriatric groups included 12, 12, 12, 24 and 20 dogs, respectively, with equal numbers of males and females. The intervention study comprised 24 young laboratory Beagles assigned to control, rapamycin, canagliflozin or dietary-restriction groups for 3 months (n = 6 per group; three males and three females). Body weight, complete blood count and a 40-target serum cytokine array were measured

A separate 90-day intervention cohort comprised 24 young intact Beagles assigned to control, rapamycin, canagliflozin or dietary-restriction groups (n = 6 per group). All 24 animals contributed CBC measurements. Serum cytokine profiling included five rapamycin-treated dogs and six dogs from each remaining group because one rapamycin serum sample was unavailable.

### Age-associated variation in peripheral blood cell counts

Systematic profiling showed selective changes across the CBC profile with age. After Benjamini-Hochberg adjustment, two-way ANOVA identified significant age-group effects in 9 of 17 CBC parameters, whereas no main effect of sex met the adjusted *P* value threshold (Fig. 2a,b and Supplementary Tables 1–3). Absolute leukocyte counts showed the largest age effects, including BASO# (partial ηp² = 0.300), WBC (partial η_p_² = 0.225), NEUT# (partial η_p_² = 0.208) and MONO# (partial η_p_² = 0.208). Five erythrocyte indices also varied significantly across age groups: HGB, HCT, MCV, MCHC and RDW-SD. RBC, HGB, HCT and MCHC showed significant interactions between age group and sex (Fig. 2b and Supplementary Table 3). Pairwise correlation analysis further showed coordinated and dynamic variation among the four absolute leukocyte counts, with the strongest association between WBC and NEUT# (Spearman’s ρ = 0.94; Fig. 2c).

**Fig. 2.**
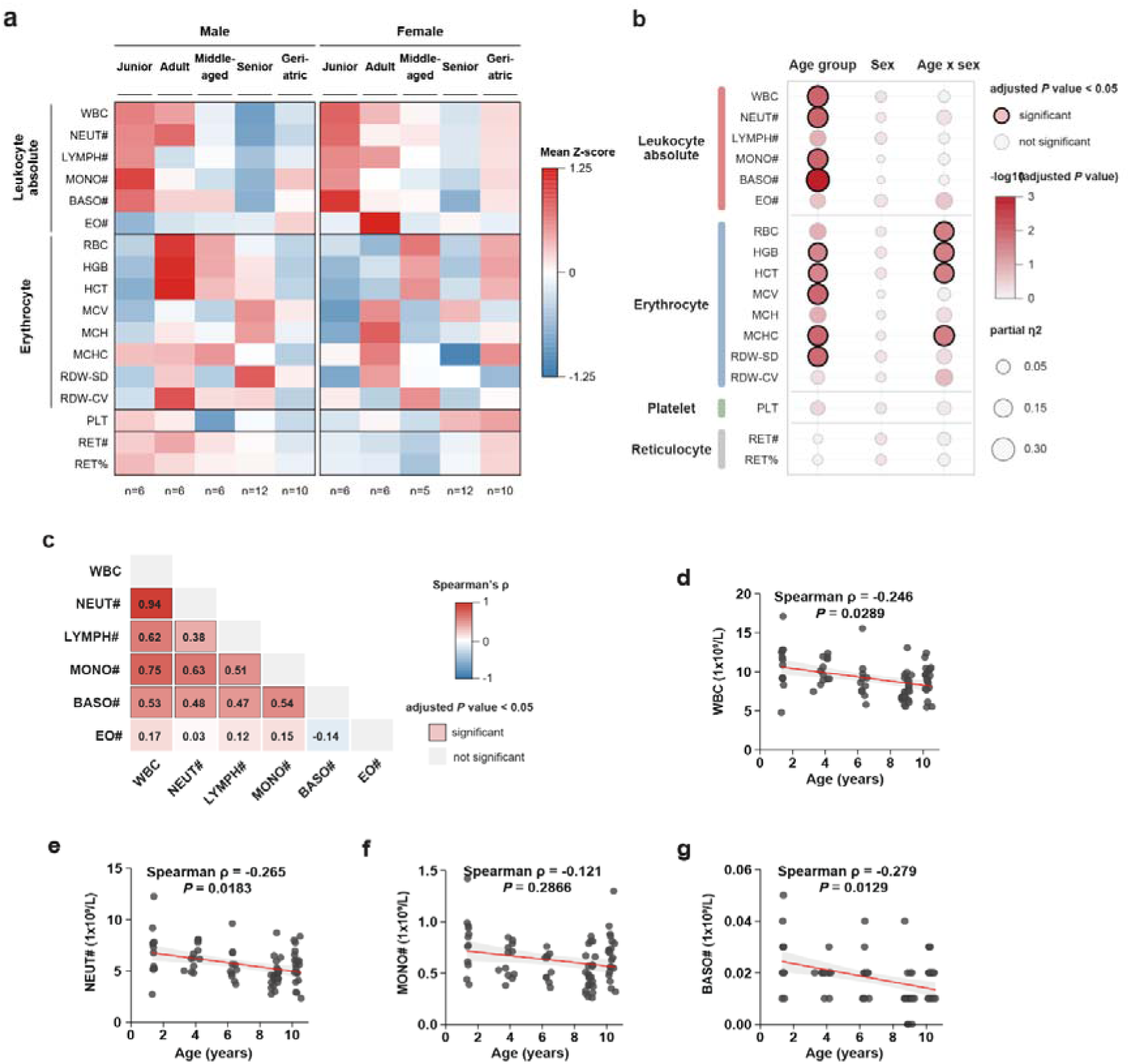
Age-related variation in canine complete blood count profiles. a, Mean Z-score heat map of complete blood count (CBC) parameters across age groups, stratified by sex. b, Effects of age group, sex and their interaction on CBC parameters, assessed by two-way analysis of variance (ANOVA). Bubble area represents partial ηp². Colour denotes -log10-transformed adjusted *P* values, and black outlines denote adjusted *P* value < 0.05. c, Pairwise Spearman correlations among absolute leukocyte counts. Values indicate Spearman’s ρ, and black outlines denote adjusted *P* value < 0.05. d,e,f,g, Associations between continuous age and WBC (d), NEUT# (e), MONO# (f) and BASO# (g). Red lines show linear fits with 95% confidence intervals as visual guides only; statistical inference was based on two-sided Spearman rank correlation. For b and c, P values were adjusted using the Benjamini-Hochberg procedure. Values shown in d,e,f,g are nominal P values. Nominal and adjusted P values for the five prespecified continuous-age correlations are provided in Supplementary Table 4. All Spearman correlation tests were two-sided. Unless otherwise indicated, n = 79 dogs. The junior, adult, middle-aged, senior and geriatric groups comprised 6, 6, 6, 12 and 10 males, and 6, 6, 5, 12 and 10 females, respectively. RET# and RET% data were available for 78 dogs owing to one missing observation

Continuous-age correlation analyses further supported these patterns. WBC, NEUT# and BASO# were negatively associated with age and remained significant after Benjamini-Hochberg adjustment across the five prespecified outcomes, while LYMPH# and MONO# were not (Fig. 2d, e, f, g). Complete results are provided in Supplementary Table 4. When analysed across age groups, all four counts were generally lower in senior dogs than in younger groups, but several increased again in geriatric dogs (Supplementary Fig. 1). Although none met the adjusted *P* value threshold for a main effect of sex or interaction between age group and sex, sex-stratified one-way ANOVA identified significant age-group contrasts more frequently in males than in females (Supplementary Fig. 1).

### Serum cytokine profiles across age groups

To evaluate systemic inflammatory and regulatory markers, we examined serum cytokine profiles. Mean Z-score heatmaps of 40 cytokines showed distinct patterns in males and females (Fig. 3a). Hierarchical clustering grouped the cytokines into three modules with different centroid trajectories across age groups (Fig. 3a, b).

**Fig. 3.**
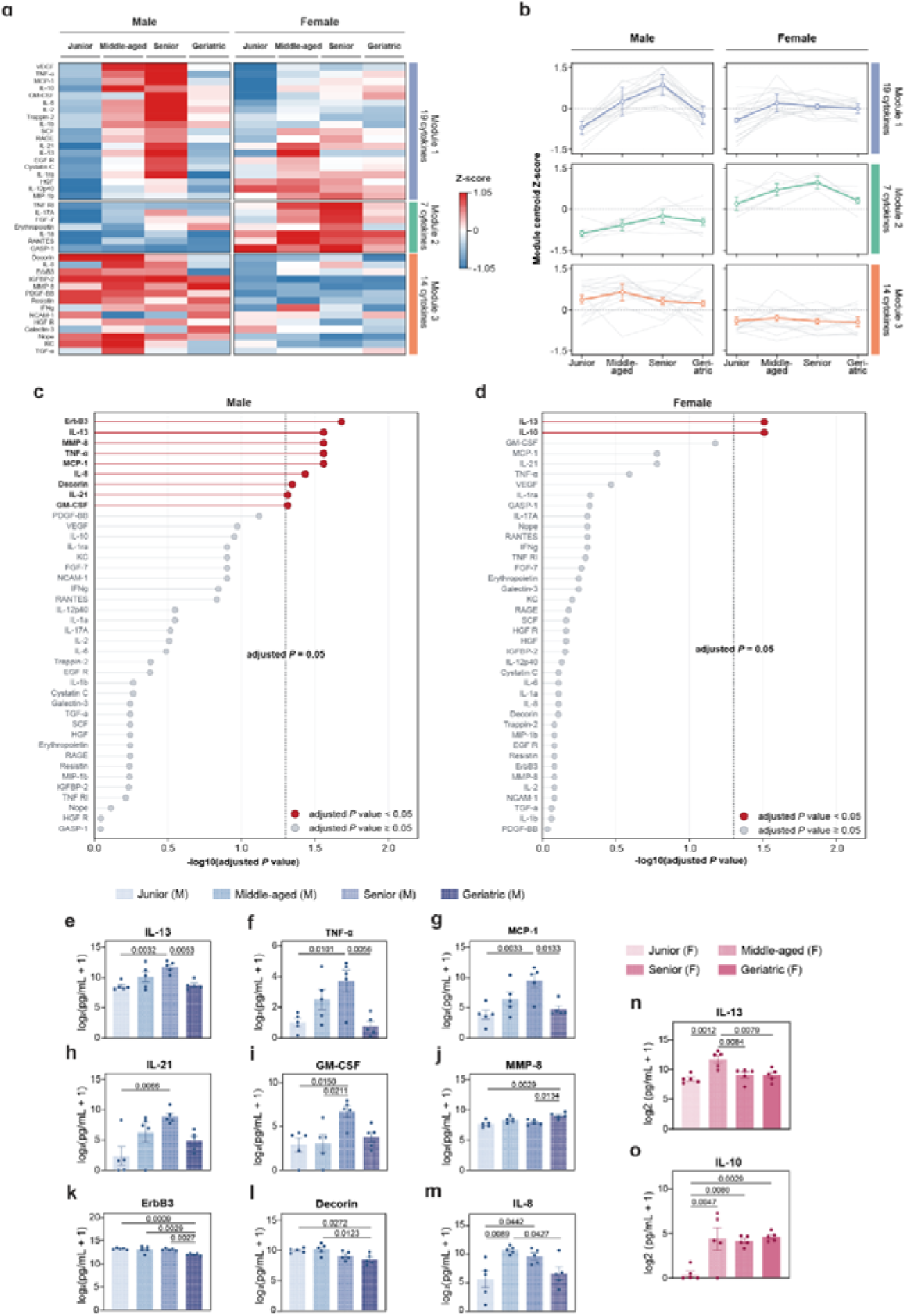
Age-associated variation in sex-stratified serum cytokine profiles. a, Mean Z-score heat map of 40 cytokines across age groups, stratified by sex. Cytokines were grouped into three modules by Ward hierarchical clustering. b, Age-group trajectories of module-centroid Z-scores. Coloured lines show mean ± s.e.m., and grey lines show the mean trajectories of individual cytokines. c,d, Age-group effects on cytokine concentrations in males (c) and females (d), assessed by one-way ANOVA. Values show -log10-transformed adjusted *P* values, and red points indicate adjusted *P* value < 0.05. e-m,n,o, Representative cytokine concentrations across age groups in males (e-m) and females (n,o), respectively. Bars show mean ± s.e.m., and points represent individual dogs. Comparisons used one-way ANOVA followed by Tukey’s multiple-comparisons test. Values above brackets are Tukey-adjusted *P* values. Cytokine concentrations were analysed on the log2(pg/mL + 1) scale. In c and d, *P* values were adjusted separately within each sex using the Benjamini-Hochberg procedure.

Sex-stratified analyses identified more age-associated cytokines in males than in females, but this difference did not establish that the age effects differed statistically between sexes. After Benjamini-Hochberg adjustment within each sex, nine cytokines differed significantly across male age groups: ErbB3, IL-13, MMP-8, TNF-α, MCP-1, IL-8, Decorin, IL-21 and GM-CSF (Fig. 3c). In females, only IL-13 and IL-10 met the same threshold (Fig. 3d). However, this unequal number of significant features does not inherently establish that aging trajectories differ statistically between the sexes. To test this directly, we performed a formal two-way ANOVA, which identified nominal age-group-by-sex interactions for IL-13, TNF-α, IL-21, MMP-8, ErbB3, and GM-CSF, but none remained significant after FDR correction (Supplementary Figure 2).

Individual-level distributions further resolved these cytokine-specific and stage-specific patterns. In males, IL-13, TNF-α, MCP-1, IL-21 and GM-CSF were higher in senior than in junior dogs but declined in the geriatric group (Fig. 3e-i). IL-8 was highest in the middle-aged group, whereas MMP-8 was highest in the geriatric group (Fig. 3j,m). Conversely, ErbB3 and Decorin declined towards the geriatric group (Fig. 3k,l). In females, IL-13 was highest in the middle-aged group, whereas IL-10 was higher in every older group than in the junior group (Fig. 3n,o). Overall, these results highlight heterogeneous, sex-stratified cytokine patterns across age groups, rather than uniform inflammatory trajectories.

### Exploratory intervention-associated differences in body weight and leukocyte endpoints

Body-weight trajectories differed descriptively among the three interventions over 90 days (Fig. 4a-c). Rapamycin increased body weight in both sexes (Fig. 4a). During canagliflozin treatment, body weight began to decline in males during the first month and in females during the second month; this descriptive difference was not tested using a treatment-by-sex interaction (Fig. 4b). Dietary restriction reduced body weight, with a larger descriptive decline in males (Fig. 4c).

**Fig. 4.**
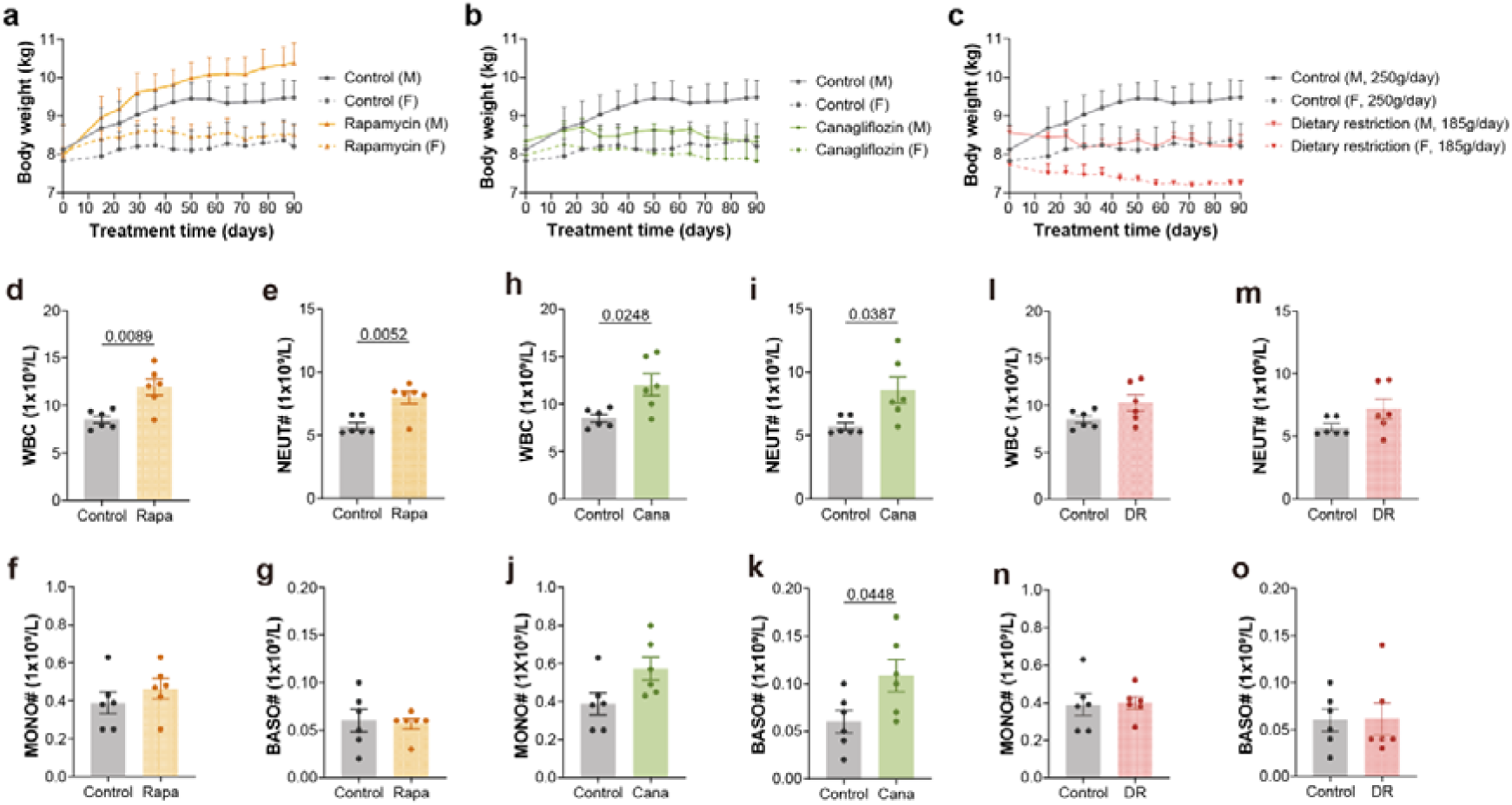
Effects of short-term rapamycin, canagliflozin and dietary restriction on body weight and leukocyte endpoints. a-c, Body-weight trajectories during 90 days of rapamycin (a), canagliflozin (b) or dietary restriction (c), stratified by sex. Lines show mean ± s.e.m. Daily food allowances were 250 g in the control, rapamycin and canagliflozin groups and 185 g in the dietary-restriction group. d-g,h-k,l-o, WBC, NEUT#, MONO# and BASO# counts following rapamycin (d-g), canagliflozin (h-k) and dietary restriction (l-o), respectively. Bars show mean ± s.e.m., and points represent individual dogs. Comparisons were made using two-sided unpaired Welch’s t-tests. Values above brackets are nominal P values. Complete nominal and adjusted P values are provided in Supplementary Table 5. For a-c, n = 3 dogs per sex within each group. For d-o, n = 6 dogs per group. Rapa, rapamycin; Cana, canagliflozin; DR, dietary restriction. Body-weight trajectories are descriptive; no treatment-by-sex interaction was tested.

At the study endpoint, WBC, NEUT#, MONO# and BASO# were compared with concurrent controls because these measures varied across age groups. Based on nominal P values (Fig. 4), the rapamycin group had higher WBC and NEUT# counts (both *P* < 0.01), and the canagliflozin group had higher WBC, NEUT# and BASO# counts (all *P* < 0.05). Dietary restriction showed no nominal differences in the four leukocyte measures. However, after Benjamini-Hochberg adjustment across all 12 comparisons, none remained significant at FDR < 0.05; the lowest adjusted *P* value was 0.0536 for WBC and NEUT# in the rapamycin group. Complete results are provided in Supplementary Table 5.

### Exploratory intervention-associated differences in serum cytokine profiles

At the 90-day endpoint, serum cytokine profiles showed exploratory intervention-associated differences (Fig. 5). Principal-component analysis showed clear separation between the rapamycin and control groups, partial separation following canagliflozin treatment, and substantial overlap between the dietary-restriction and control groups (Fig. 5a,k,p).

**Fig. 5.**
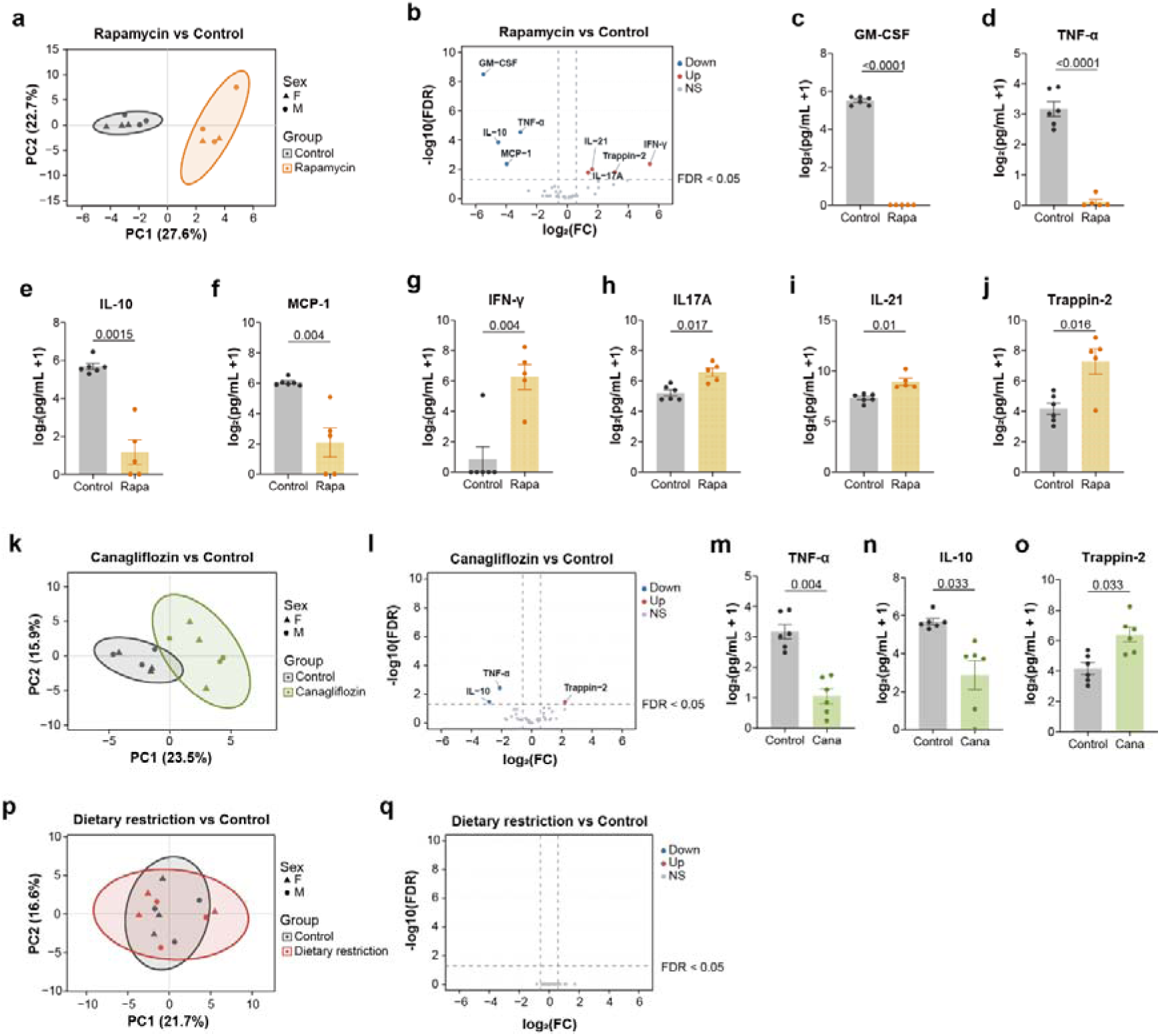
Short-term intervention-associated differences in serum cytokine profiles. a,k,p, Principal-component analysis (PCA) of serum cytokine profiles following rapamycin (a), canagliflozin (k) and dietary restriction (p). Cytokine concentrations were log2-transformed, and each cytokine was scaled before PCA. Points represent individual dogs, point shapes indicate sex, and ellipses show 95% data ellipses. b, l, q, Volcano plots of differential serum cytokine abundance following rapamycin (b), canagliflozin (l) and dietary restriction (q). Differential abundance was assessed using limma-style moderated two-sided t-tests. Raw *P* values were adjusted across the 40 cytokines within each comparison using the Benjamini–Hochberg procedure to control the false discovery rate (FDR). Coloured points indicate FDR < 0.05 and an absolute fold change of at least 1.5. c–j, m–o, Bar plots of cytokines meeting these thresholds following rapamycin (c–j) or canagliflozin treatment (m–o). Bars show mean ± s.e.m.; points represent individual dogs, and values above brackets are Benjamini-Hochberg-adjusted *P* values from the differential-abundance analyses. For a-j, the rapamycin and control groups included n = 5 and n = 6 dogs, respectively, after exclusion of one rapamycin serum sample. For k-o and p-q, n = 6 dogs per group. Rapa, rapamycin; Cana, canagliflozin; DR, dietary restriction. Cytokines were measured only at the 90-day endpoint; these exploratory analyses estimate between-group differences rather than within-individual changes from baseline.

At an adjusted *P* value threshold of 0.05 and an absolute fold-change threshold of 1.5, eight cytokines differed between the rapamycin and control groups. The rapamycin group had lower GM-CSF, IL-10, MCP-1 and TNF-α but higher IFN-γ, IL-17A, IL-21 and Trappin-2 (Fig. 5b-j). The canagliflozin group had lower IL-10 and TNF-α and higher Trappin-2 than the control group (Fig. 5l-o). No cytokines passed the predefined thresholds following dietary restriction (Fig. 5q).

Among the three interventions, rapamycin was associated with the largest number of differential cytokines. Canagliflozin showed fewer cytokine differences and descriptive sex-stratified body-weight trajectories, whereas dietary restriction reduced body weight without detectable differences in the measured leukocyte or cytokine endpoints. Because cytokines were measured only at the endpoint, these findings represent between-group differences and do not establish within-individual changes from baseline.

## Discussion

Aging is closely linked to immune dysfunction and chronic inflammation (Ferrucci and Fabbri 2018; Franceschi et al. 2000; Franceschi et al. 2018; López-Otín et al. 2023). In this controlled cohort of laboratory Beagles, peripheral blood cell counts and serum cytokine profiles showed heterogeneous, non-linear age-associated patterns. Several leukocyte counts declined with continuous age but increased again in geriatric dogs, whereas cytokines changed in different directions across age groups. These observations indicate that canine immune aging is better described as stage-specific patterns than as a uniform decline or a steady increase in inflammation. The cytokine findings are described as sex-stratified patterns rather than statistically established sex-dimorphic trajectories because no interaction remained significant after FDR correction. Standardized breed, diet, housing, and intact gonadal status reduced several sources of variation and provided a controlled complement to more heterogeneous companion-dog cohorts (Kaeberlein et al. 2016; McMahon et al. 2025; Schmid et al. 2024).

Absolute leukocyte counts showed the clearest age-associated changes among the CBC parameters. WBC, NEUT# and BASO# remained negatively associated with continuous age after correction across five outcomes, whereas LYMPH# and MONO# did not. Across age groups, all four counts were generally lower in senior dogs than in younger groups, but several increased again in geriatric dogs. Similar age-associated changes in circulating immune cells have been reported in previous canine studies (Alexander et al. 2018; Jiménez 2023; McMahon et al. 2025; Schmid et al. 2024). However, the late-life increase in this study should be interpreted cautiously. Because the cohort was cross-sectional, it cannot determine whether this pattern reflects within-individual change, selective survival, or differences in age-group composition. Sex effects also differed among hematologic endpoints. None of the four leukocyte counts met the adjusted *P* value threshold for a main effect of sex or age-by-sex interaction, whereas RBC, HGB, HCT and MCHC showed significant age-by-sex interactions. Sex-related hematologic aging in this cohort therefore appeared to be endpoint-specific rather than shared across the entire CBC profile.

Serum cytokine profiles were more heterogeneous than leukocyte counts and showed different descriptive patterns in males and females. In males, IL-13, TNF-α, MCP-1, IL-21 and GM-CSF peaked in senior dogs before declining in the geriatric group. IL-8 peaked in middle-aged males and MMP-8 in geriatric males, whereas ErbB3 and Decorin declined towards the geriatric group. In females, IL-13 was highest in the middle-aged group, whereas IL-10 was higher in every older group than in the junior group. These non-monotonic patterns resemble molecular changes reported across the human lifespan and indicate that inflammatory, regulatory and tissue-remodeling signals do not change in parallel (Lehallier et al. 2019; Shen et al. 2024). More cytokines met the age-group significance threshold in males than in females. However, unequal numbers of significant results do not demonstrate that age effects differ between sexes, and no formal cytokine interaction remained significant after FDR correction. Estrous stage was not recorded, although none of the female dogs was pregnant, lactating or being used for breeding at sampling. Unmeasured variation across the estrous cycle may therefore have contributed to variability in the sex-stratified cytokine results. Additionally, owing to assay capacity constraints, the adult age group was omitted from the cytokine profiling. This omission limits our resolution between the junior and middle-aged stages and prevents the estimation of fully continuous lifespan trajectories. Larger longitudinal studies with adequately powered interaction tests are needed to determine whether these patterns represent robust sex-dependent immune aging.

Rapamycin was associated with the broadest endpoint immune response among the three interventions. At the 90-day endpoint, the rapamycin group had nominally higher WBC and NEUT# counts than the control group, but neither comparison remained significant after correction across the 12 CBC tests. Rapamycin was also associated with lower GM-CSF, IL-10, MCP-1 and TNF-α, but higher IFN-γ, IL-17A, IL-21 and Trappin-2. This mixed cytokine response does not support a simple anti-inflammatory interpretation and instead points to broader immune modulation. Several rapamycin-responsive measures were also associated with age in the cross-sectional analysis. However, this overlap does not show that rapamycin reverses immune aging because the intervention was conducted in young dogs. Previous canine rapamycin studies have focused mainly on safety and cardiac outcomes (Barnett et al. 2023; Urfer et al. 2017), and the ongoing TRIAD trial will assess longer-term effects in a larger cohort of middle-aged dogs (Coleman et al. 2025). This exploratory study adds early peripheral immune endpoints to this developing evidence base.

Canagliflozin was associated with a narrower endpoint response than rapamycin. At the endpoint, the canagliflozin group had nominally higher WBC, NEUT# and BASO# counts, but none remained significant after correction across the 12 CBC tests. The group also had lower TNF-α and IL-10 and higher Trappin-2 than the control group. Descriptive body-weight trajectories showed an earlier and larger reduction in males than in females. These observations are consistent with mouse studies in which canagliflozin extended median lifespan in males but not females and produced sex-dependent metabolic and neuroinflammatory effects (Jayarathne et al. 2022; Miller et al. 2020). They motivate further study of possible sex-dependent intervention responses. However, each intervention group contained only three dogs of each sex, and body-weight trajectories were analyzed descriptively. We therefore did not perform sex-stratified immune analyses or formal treatment-by-sex interaction tests. The apparent difference in body-weight response should be regarded as hypothesis-generating rather than evidence of a sex-specific treatment effect in dogs. Importantly, while blood drug concentrations and routine clinical chemistry indices confirmed systemic exposure and general clinical tolerability for both the rapamycin and canagliflozin regimens, comprehensive pharmacokinetic, pharmacodynamic, and metabolic profiling are warranted in the future studies.

Dietary restriction reduced body weight but did not significantly alter the measured leukocyte counts or serum cytokines over 90 days. This result should not be interpreted as evidence against dietary restriction as a geroscience intervention. Long-term 25% dietary restriction extended median lifespan and delayed age-related disease in Labrador retrievers (Kealy et al. 2002; Lawler et al. 2008), whereas two years of calorie restriction altered thymic function and circulating immune mediators in humans (Spadaro et al. 2022). The absence of a detectable immune response in the present study may reflect the shorter exposure, the use of young clinically healthy dogs, the small intervention groups, or the specific endpoints examined. In addition, cytokines were measured only at the 90-day endpoint. The intervention findings therefore represent between-group endpoint differences rather than within-individual changes from baseline. Longer studies with baseline measurements will be needed to determine whether dietary restriction produces delayed or age-dependent immune effects.

Overall, our study describes stage-specific hematologic changes and sex-stratified cytokine patterns across the canine lifespan, together with exploratory short-term responses to three candidate interventions. Several erythrocyte indices showed significant age-group-by-sex interactions, but no cytokine interaction remained significant after FDR correction. Thus, these data do not establish sex-dimorphic cytokine aging. The use of intact laboratory Beagles reduced one source of endocrine heterogeneity, although unmeasured estrous stage and the standardized colony setting limit interpretation and generalization. Rapamycin and canagliflozin were associated with endpoint immune differences, whereas short-term dietary restriction did not alter the measured endpoints. These intervention findings are exploratory and larger longitudinal studies with adequately powered sex-specific designs are needed to validate these early-life signatures and translate them into actionable geroscience.

## Acknowledgments

We thank the team at Anling Biomedical (Shenzhen) Co., Ltd. for assistance with the canine experiments.

## Author contributions

Meiling Lai: Experiment validation, Data curation, Writing - original draft, review and editing. Fengge Xu, Yingxia Xu and Albert Mironenkov: Experiment validation and Data curation. Yi Jin: Resource arrangement and Data curation. Shengxiang Zhang and Jianhong Pan: Investigation and technical guidance on experimental procedures. Ming Li: Funding acquisition, Writing - review and editing. Baichuan Deng: Supervision, Writing - review and editing. Jian-Kang Zhu: Funding acquisition, Supervision, Writing - review and editing. Yu-Xuan Lyu: Conceptualization, Funding acquisition, Supervision, Writing - review and editing.

## Ethics approval and animal welfare

All animal procedures were approved by the Institutional Animal Care and Use Committee of Anling Biomedical (Shenzhen) Co., Ltd. (approval no. 2025-02-ALSHZH-21). All procedures complied with applicable institutional and national guidelines, the NIH Guide for the Care and Use of Laboratory Animals, and the ARRIVE guidelines.

## Data availability

The source data supporting the findings of this study are provided in the accompanying Source Data file. The complete nominal and Benjamini-Hochberg-adjusted *P* values for the five prespecified continuous-age correlations and the 12 intervention CBC comparisons are reported in Supplementary Tables 4 and 5, respectively. Additional data are available from the corresponding author upon reasonable request.

